# Use of nanomaterials-modified carbon microfibre electrode material for superior electrochemical performance in lake sediment inoculated microbial fuel cells

**DOI:** 10.1101/2023.07.10.548441

**Authors:** Maheshi Somasiri, Tanusha Amandani, Charitha Basnayaka, Ahmed Ahsan, Gayani P Dilangani, Ajith C. Herath, Sampath Bandara, Zumaira Nazeer, Nirath Thilini, Godfrey Kyazze, Eustace Y. Fernando

## Abstract

High cathodic overpotential of the oxygen reduction reaction (ORR) in MFC carbon-based cathodes is one of the key barriers to the widespread adoption of the technology. Current Pt-based ORR catalysts are expensive. The use of novel and inexpensive catalysts as replacements for platinum is therefore desirable. In this study, nanomaterials were directly chemically synthesized on carbon microfiber electrodes to improve the performance of lake sediment inoculated MFCs. Nanomaterial of MnO_2_, MnO_2_/polyaniline (PANI), ZnO/NiO and ZnO/NiO/PANI attachments were directly chemically synthesized on the carbon material and used as cathode electrodes. The maximum power densities recorded for the different treatments were; MnO_2_ 78.5 mW/m^2^, MnO_2_/PANI (Polyaniline) 141.6 mW/m^2^, ZnO/NiO 67.6 mW/m^2^, and ZnO/NiO/PANI 129.4 mW/m^2^. The current and poswer densities were more than six-fold higher in ZnO/NiO/PANI and MnO_2_/PANI nanoparticle modified cathodes compared to the control MFCs with no catalyst. Cyclic voltammetry (CV) and FTIR data and SEM images suggest that the nanoparticle attached carbon material is morphologically, chemically and electrochemically different from the controls with no nanomaterial attachment. The outcome of this study demonstrates that nanomaterials-incorporated carbon microfiber cathodes bring about significant enhancements to power densities and may potentially have applications in cost-effective MFCs.

## 1. Introduction

The energy needs cannot be sustained solely by fossil fuels at the end of the 21^st^ century as they are not substantial enough and pose environmental problems such as contributing to global warming and climate change. As a result, there is a pressing need for renewable alternative energy generation. Microbial fuel cells (MFCs) have emerged in recent years as a promising technology for renewable alternative energy and wastewater treatment.

In 1911, English botanist M. C. Potter (Potter, 1911) discovered that when a platinum electrode was put into a liquid solution of yeast and *Escherichia coli*, bacteria could create an electric current. This finding marked the beginning of Microbial Fuel Cell (MFC) research and the subsequent technical advancements. Simply, MFCs employ microorganisms as biocatalysts to oxidize organic materials and transmit electrons to the anodic surface via substrate oxidation for bioelectricity production (Logan, 2009). Almost any source of biodegradable organic matter, including basic molecules like carbohydrates and proteins, as well as complex combinations of organic matter, found in human, animal, and food-processing waste streams, may be employed in MFC for power production. The MFC is an attractive technology for renewable bioelectricity generation from biomass due to microorganisms’ ability to utilize a variety of fuels (Logan, 2009).

A standard MFC is made up of two chambers separated by a Proton Exchange Membrane (PEM), an anode, and a cathode. Electrons are delivered from the anode compartment to the cathode compartment through an external circuit, where they couple with protons and oxygen to produce water via the procedure described below (Ucar et al., 2017).

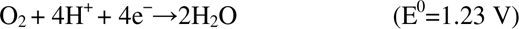

Electrons generated by anodic biofilm bacteria following oxidation of an electron source (usually an organic substrate) are initially transported to the anode under anoxic conditions in a conventional MFC setup. This suggests that the exo-electrogenic biofilm bacteria employ the anode as an electron acceptor for anaerobic respiration(Li et al., 2018).

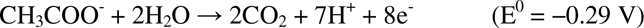

Protons produced during the oxidation process typically move to the cathode via a PEM, which inhibits oxygen passage into the anodic chamber. Electrons are sent to the cathode through an external circuit with a resistance load. Water is formed when electrons, protons, and dissolved oxygen in the catholyte react on the cathode surface (Li et al., 2018).

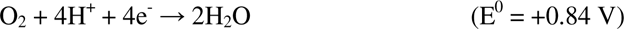

Although an MFC’s power density remains low when compared to a hydrogen fuel cell, its renewable and widely available fuel sources, as well as its moderate operational conditions, make it highly promising in renewable energy generation, wastewater treatment, and as portable power supply or as a continuous power sources for remote sensing electronic devices.

MFC microorganisms are typically bacteria that can degrade organic matter via anaerobic oxidation of organic substrates. The bacteria consume organic matter and use it as a source of energy in this process, producing electrons as a byproduct. These electrons can be captured and used to power a generator. Bacteria and archaea make up the microbial community in MFCs, with bacteria being the most common. Although, methanogenic archaea can pose a detrimental effect on MFC performance by diverting a portion of the available pool of electrons in organic substrates into methanogenic pathways (Jayathillake et al., 2022). Based on their method of transferring electrons to the anode, the microorganisms used in MFCs can be divided into two groups: Direct electron transfer microorganisms and indirect (mediated) electron transfer. Electrogenic bacteria, such as *Shewanella* and *Geobacter*, can directly transfer electrons to the anode surface. Genera such as *Pseudomonas, Rhodopseudomonas*, and *Clostridium* are involved in extracellular electron transfer (EET) pathways and some do not directly transfer electrons to the anode (Konovalova et al., 2018).

Electric current is known to be generated in MFCs by exo-electrogenic bacteria such as *Shewanella* and *Geobacter*. Gram-negative bacteria *Shewanella* spp. and *Geobacter* spp. can generate electricity by using a variety of electron acceptors, including Fe (III) and Mn (IV) (Liu et al., 2004). These bacteria are electrochemically active due to specific cytochromes on the outer side of the cell membrane (Logan B., 2009). The MFC anode can function as the final electron acceptor. Some *Geobacter* and *Shewanella* strains are capable of producing electrically conducting pilli (nanowires), allowing the microorganism to use more distant solid electronic acceptors (Logan B., 2009).

In most cases, microorganisms are electrochemically inactive and cannot directly transfer electrons to the electrode, but they are involved in EET pathways. These bacteria are important for the performance of MFCs because they produce various electron shuttles, such as quinones and flavins, which help to transfer electrons to the anode. *Pseudomonas* spp. like *Pseudomonas aeruginosa* produce pyocyanin, a mediator molecule that is used as a shuttle for electron transfer (Logan B., 2009).

In addition to bacteria, other microorganisms have also been explored for use in MFCs. For example, algae have been used in MFCs to generate electricity through photosynthesis. A sediment-type self-sustained phototrophic microbial fuel cell (MFC) run by Zhen He and his colleagues in the year 2009 used a synergistic interaction between photosynthetic microorganisms and heterotrophic bacteria, convert sunlight into energy, which is then used to produce electrons that can be captured and used to generate electricity (He et al., 2009).

Identifying the limiting parameters in an MFC system is crucial for improving MFC performance further. An MFC’s power output is limited by internal resistance, which includes anode resistance, cathode resistance, electrolyte resistance, and (if present) membrane resistance (Logan et al., 2006). Internal resistance can be lowered by increasing the anode surface area, cathode surface area, proton exchange membrane surface area, electrolyte ionic strength, or pH (Oh & Logan, 2006).

The cathodic overpotential of oxygen reduction reaction (ORR) at MFC cathode-based graphite is a limiting factor for the technology’s widespread development. The cathode performance can be improved in two ways, either by using a catalyst to reduce the activation energy or by increasing the specific surface area of the graphite material (Erable et al., 2009). Currently, available ORR catalyst materials such as platinum or palladium powder are expensive and decrease the practical applicability of MFC systems.

Many studies have indicated a boost in ORR rates using novel and inexpensive catalysts, while other researchers have employed non-catalyzed three-dimensional graphite in cathode fabrication, such as cloth, felt, sponge, or granules.

Carbon fiber fabrics enable 3D integration of electrode material in reactors, enhancing fuel mass transfer and mediator diffusion due to their porous features. Additionally, larger pores or channel diameters of 3D carbon materials allow bacterial cell penetration, generating interest among MFC anode researchers (Fan et al., 2021).

The use of nanomaterials and nanotechnology has revolutionized the fabrication of components in microbial fuel cells (MFCs), leading to an improved performance in electron conductivity, power density, cost, thermal stability, and catalysis of ORR rate.

Notably, modification of MFC electrodes (anode and cathode) with nanomaterials has a critical impact on overall MFC performance, enhancing bioenergy production. Examples of nanomaterials used for electrode modification in MFCs include metal nanoparticles and metal oxides (such as CeO_2_, TiO_2_, ZnO, SiO_2_, Al_2_O_3_, and MnO_2_) (Khoo et al., 2020).

This study was designed to test how the nanomaterial-modified carbon microfiber electrode material can enhance the performance of lake sediment inoculated microbial fuel cells by enhancing the ORR at the cathodic component of the cells. Primary objective of the study was to fabricate various nanoparticle embedded carbon microfiber electrodes by using the techniques direct electro-synthesis and electro-deposition and to conduct a comparative study into the electrochemical performance of these novel nanoparticle-embedded electrode constructs when used in two-chamber MFCs. Of particular interest was the performance of these novel materials was their capability and efficiency of catalyzing ORR reactions in MFCs.

## 2.0 Materials and Method

### 2.1 Construction of two chambered Microbial Fuel Cell

The MFC system was constructed as shown in Figure 1 using a cast acrylic sheet. Three cells were constructed, one as the control and the other two as the replicates.

**Figure 1:**
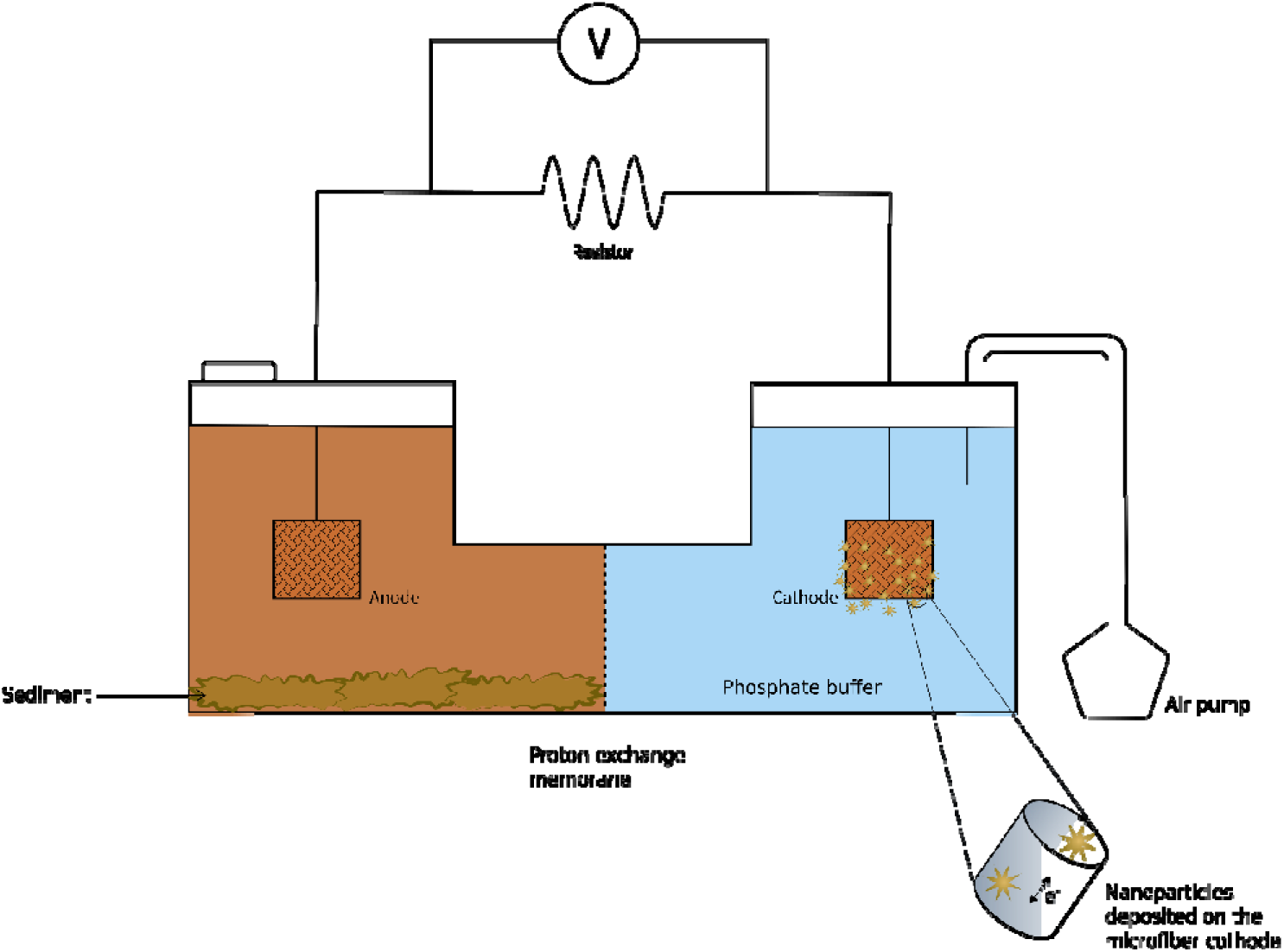
Schematic diagram of the two chamber MFC system with nanomaterials deposited woven microfiber cathode

**Figure 2:**
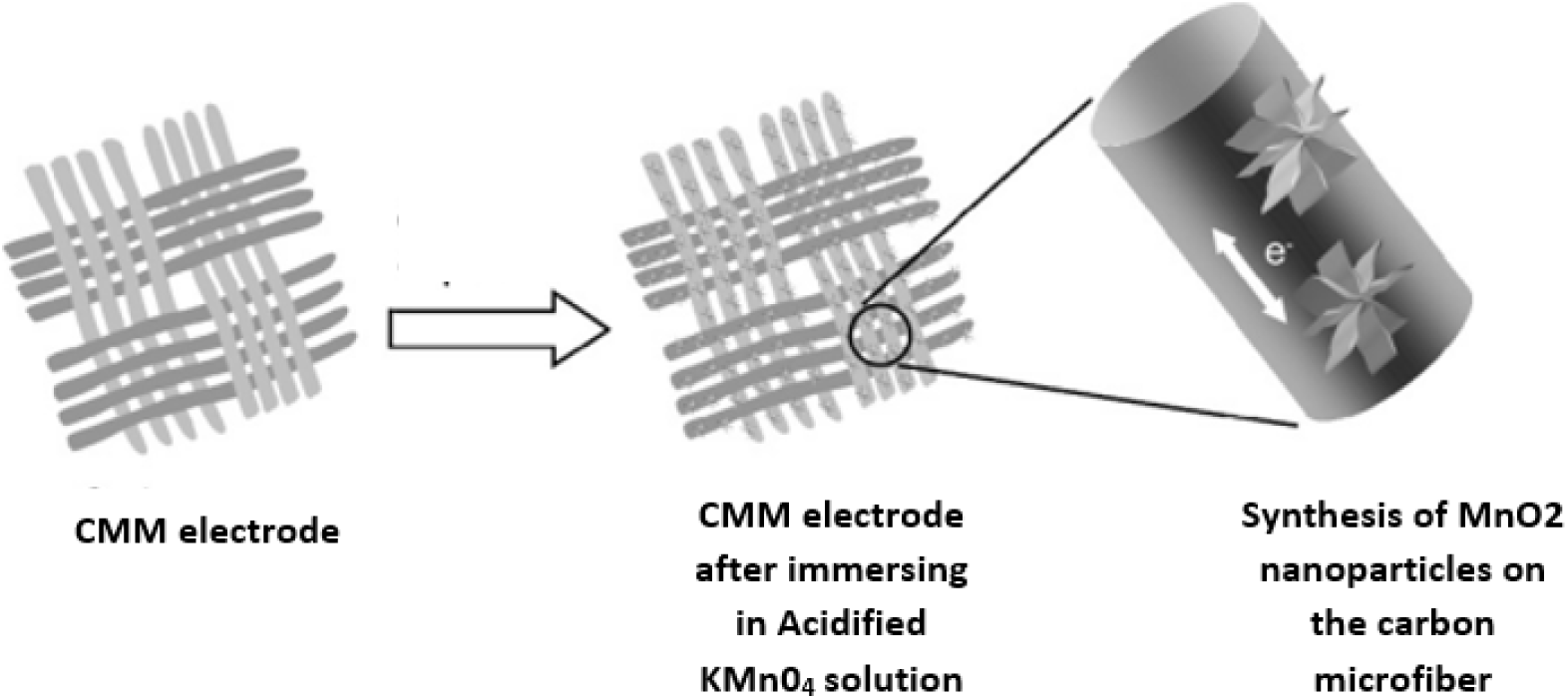
Graphical representation of direct synthesis of Manganese oxide nanoparticles on the carbon microfiber material electrode

#### 2.1.1 Anode and Cathode Chambers

Both the anode and cathode chambers were roughly 300cm^3^ in capacity and were separated by a Nafion 212 Proton Exchange membrane. To maintain the anode chamber anaerobically, it was sealed with an airtight sealer. To maintain the cathode chamber aerated, oxygen was supplied by an air sparger at a constant flow rate.

To maintain the cells aerated, oxygen was delivered to the cathode chambers of all cells at first using an air sparger with a constant flow rate.

#### 2.1.2 Pretreatment of Nafion 212 proton exchange membrane

The Nafion 212 proton exchange membrane was first immersed in 3% H_2_O_2_, then in deionized or distilled water for two hours (2 hours), and then in 0.5M sulfuric acid for another hour. All the solutions were approximately 80°C (Figure), and the membrane was rinsed in deionized water before and after each step. To keep it from shriveling, it was kept in deionized water until it was put into the cell.

#### 2.1.3 Electrodes

Electrodes were connected externally by insulated copper wire, with resistors (10 Ω to 10,000 Ω) in-between. A programmable data logging Voltmeter (PicoLog™ 1012, Pico Technology, UK) was attached parallelly to the respective resistor.

#### 2.1.4 Media of each chamber

In the anode chamber, 250cm^3^ of Semi defined minimal media (Glucose/ acetate as carbon source) was added. Semi-defined media will contain Glucose or acetate 2g /L, NH_4_Cl 0.46 g/L, yeast extract 0.1 g/L, peptone 0.5 g/L, K_2_HPO_4_ 5.05 g/L, KH_2_PO_4_ 2.84 g/L.

In the cathode chamber, 50 mmol/dm^3^ phosphate buffer with pH 7 (K_2_HPO_4_ 5.05 g/L and KH_2_PO_4_ 2.84 g/ L) was used.

#### 2.1.5 Sample collection

Pankulama Lake (8.6264915,81.0407939), Morawawa, North Central Province, Sri Lanka was used to gather lake sediment samples. Water-logged locations with a dark/black color sediments due to the presence of pyrite deposition and a strong sulfide smell were carefully chosen for sampling because they are a good source of exoelectrogens (especially those that can reduce manganese and iron oxides) (Nazeer Z. & Fernando, 2022). Sediment samples were collected into 50 ml Sample collecting tubes and immediately sealed and kept at room temperature until they were used in MFC experiments.

#### 2.1.6 Microbial inoculum

Approximately 250 cm^3^ of semi-defined minimal media with a glucose solution of 2g/L was added to the anode chamber. Glucose 2g/L, NH_4_Cl 0.46 g/L, Yeast extract 0.1 g/L, Peptone 0.5 g/L, K_2_HPO_4_ 5.05 g/L, KH_2_PO_4_ 2.84 g/L make up the anode solution. The salt solution and glucose solution were autoclaved separately for 20 minutes at 121°C and 15psi and added later in the anode compartment. The anode compartment was sealed airtight with silicon glue after the addition of approximately 5g of sediment sample and 200cm^3^ of semi-defined media to maintain an anaerobic environment for exoelectrogens.

#### 3.1.7 Batch operation of the MFC

After filling the anode chamber with sediment and the semi-defined minimal medium, it was operated in batch mode with intermittent substrate feeding. After 2-3 days, when the output voltages had stabilized, approximately 30% of the anode volume was replenished with fresh semi-defined media containing glucose as the exogenous feeding substrate.

The final glucose concentration at the beginning of each batch cycle was 2 g/L.

### 2.2. Preparation and modification of Electrodes

The carbon microfiber material (CMM) was obtained from Carbon Energy WT (Taiwan) as a custom-fabricated woven carbon material for electrode construction. The carbon microfiber material had an average fiber diameter of 8 ± 0.5 µm as per the manufacturer data. Each woven carbon microfiber electrode was approximately 4mm thick and 5 cm X 5 cm width and length dimensions (Jayathilake et al., 2022).

#### 2.2.1 Base Treatment of Electrodes

Base-treated CMM electrodes were prepared by submerging a 5x5 cm CMM electrode into KOH (3M) solutions and heating at 85^0^ C for six hours (6h). After the solutions cooled down to room temperature, the CMM electrode was soaked in an alkaline solution for twenty-four hours (24h) and was rinsed in deionized water three times. Finally, the CMM electrode was air-dried for twelve hours (12h) (Wang et al., 2013).

#### 2.2.2 Acid Treatment of Electrodes

Acid-treated CMM electrodes were prepared by submerging a piece of CMM electrode into HNO_3_ (5.6M) solutions and heating at 85^0^ C for six hours (6h). After the solutions cooled down to room temperature, CMM was soaked in an acidic solution for twenty-four hours (24h) and rinsed in deionized water three times. Finally, the CMM electrode was dried in a vacuum drier for twelve hours (12h) (Wang et al., 2013)

#### 2.2.3 Nanomaterial modification of Electrodes

##### 2.2.3.1 Pre-treatment of the electrode material

The untreated carbon microfiber electrode material was cut into 5 cmx5 cm dimensions were dipped in a 1M sulfuric acid solution and sonicated for 15 min. It was then sonicated again after dipping it in an acetone solution for 15 minutes. And finally, it was sonicated again in distilled water for 15 minutes. It was then air dried for twenty-four hours (24h).

##### 2.2.3.2 Electro deposition of Polyaniline (PANI) on the electrodes

To prepare PANI, 0.1 M double distilled aniline was added dropwise to 100 ml of 1M HCl until it reached a of pH 1.5. The deep green color of polyaniline was formed by a chemical oxidation process.

Electrochemical study was conducted using Scanning Electrochemical Microscope (CH instruments (Austin, TX, USA). Cyclic voltammetry Programme with a standard three electrode configurations. Modified CMM electrode (m-CMM) (2.2.3.1) (working area 5 cm × 5 cm) was used as the working electrode, platinum (Pt) was used as a counter electrode, with an Ag|AgCl (KCl 0.1molar) reference electrode completing the cell assembly. Above mentioned electrodes were placed horizontally in 10 mL glass container with formed PANI and connected them with Scanning Electrochemical Microscope instrument by connecting appropriate connecting wires. Then cyclic voltammetry Programme was chosen from the software for the run. After the m-CMM electrode became coated with PANI it was washed with distilled water and air-dried for 24 hours at ambient temperature.

##### 2.2.3.3 Direct synthesis of Manganese oxide (MnO_2_) nanoparticles on the CMM electrodes

A 100ml solution of equal volumes of 0.1M Sodium Sulphate (Na_2_SO_4_) and 0.1M Potassium permanganate (KMnO_4_) was prepared. m-CMM electrode (<u>3.2.3.1</u>) was dipped in this solution for 6 hours. Then it was kept at room temperature until it was completely dry. After deposition, this electrode was washed with distilled water to eliminate any loosely attached chemical substances.

##### 2.2.3.4 Deposition of MnO_2_ nanoparticles on a PANI-deposited CMM electrode

An m-CMM electrode (2.2.3.1) which was treated with PANI as mentioned in section 2.2.3.2 was treated with direct deposition of MnO_2_ nanoparticles as mentioned in section 2.2.3.3.

##### 2.2.3.5 Direct deposition of Zinc oxide (ZnO) nanoparticles on m-CMM electrodes

m-CMM electrode was dipped in a 100ml of equal volumes of 3 mM Zinc sulfate (ZnSO_4_.7H2O), 7mM Urea (CO(NHL)L) and Nickel sulfate (NiSOL(HLO)L) solution in a crucible. It was kept under 180 ^0^C for 2 hours and then under 360 ^0^C for 1 hour. It was then washed with distilled water to eliminate any loosely attached chemical substances.

##### 2.2.3.6 Direct deposition of Zinc oxide (ZnO) and Nickel oxide (NiO) nanoparticles on m-CMM/PANI electrodes

m-CMM/PANI electrode was dipped in a 100ml of equal volumes of 3 mM Zinc sulfate (ZnSO4.7H2O), 7mM Urea (CO(NHL)L) and Nickel sulfate (NiSOL(HLO)L) solution in a crucible. It was kept under 180 ^0^C for 2 hours and then under 360 ^0^C for 1 hour. It was then washed with distilled water to eliminate any loosely attached chemical substances.

### 2.3 MFC performance characterization

The voltage across the resistance was measured using a PicoLog 1020 (Pico Technology, UK) data logging system. Different resistances were connected ranging from 22Ω to 10,000,000Ω and the current flowing through each resistance was calculated using the equation I= E/R (where, I – current, E – voltage and R – resistance). The power dissipated across each resistance was calculated using P=IE (where, P – Power, E – voltage and I – current). The power dissipated across the resistance was normalized to the area of the electrode (25 cm^2^). The electrochemical performance of each MFC experiment was analyzed using polarization curves and power-current plots. Control-A 5cm x 5cm Non-treated CMM electrode was used as an electrode in both the cathode and anode in the control MFC system. Additional control MFC with a cathode containing a Pt loading of 0.5 mg cm^2^ was also used for comparison.

### 2.4 Cyclic voltammetry to investigate the performance of electrodes

Before initiating the CV apparatus, the counter and reference electrodes were gently washed with deionized water. The electrochemical cell was prepared by assembling the working electrode, reference electrode, and counter electrode according to the experimental setup. The working electrode is the Nanomaterial modified electrode and the reference electrode and the counter electrode are Ag|AgCl and platinum (Pt) respectively.

The phosphate buffer was used as the electrolyte and a sweep rate of 50 mV/s was used in all CV experiments. The potential range was determined by running a few cycles. Then the CV experiment was performed, and the CV curves were obtained.

### 2.5 Scanning electron microscopy (SEM) analysis of functionalized microfiber electrode surfaces

Woven carbon Microfiber material samples after chemical treatments and after introducing them to the MFC were imaged for SEM on a Carl Zeiss EVO 18 (Germany) scanning electron microscope with an optical magnification range of 20–135 ×, electron magnification range of 5x - 1,000,000 x, maximal digital zoom of 12 ×, acceleration voltage of 10 kV, equipped with secondary electron (SE) and energy dispersive X-ray spectrometer (EDS) detectors, with a nominal resolution of 10 nm or less. The microscope had a temperature-controlled sample holder (temperature range −25°C to 50°C).

### 2.6 Fourier Transform Infrared Spectroscopy (FTIR) analysis of functionalized cathodes

FTIR was used to examine chemical functionalization on pretreated electrodes (Dissanayake et al., 2021). Dried cathode fragments were extracted from acid and base pretreated electrode MFC experiments and examined with an FTIR spectrometer (Shamadzu, UK, National Institute for Fundamental Studies, Sri Lanka). Fully dried samples were put on the instrument analysis pedestal, and transmittance spectra in the mid-IR region (400cm^-1^ - 4000cm^-1^) were acquired (Nazeer & Fernando, 2022).

## 3.0 Results and Discussion

### 3.1 MFC performance characterization

Polarization curves were obtained by recording the voltage via varying the external resistance from 22 Ω to 1 MΩ when MFC voltages across the set external resistance stabilized at the maximum steady value after substrate feeding cycles. Fig. 3 (B) indicates the power density of each experiment incorporating different nanomaterials in the cathode, acid/alkali pretreated cathodes and the controls (non-treated electrode and Pt loaded). Polarization curves of all the treatments indicated notable enhancements to the electrochemical performances of the MFC systems in terms of maximum power density (P_max_) and current density (J_max_) values, when compared to the non-treated control MFC systems. The maximum current, power production, and open circuit voltage obtained in each MFC with respective treatments are given in Table 1.

**Figure 3:**
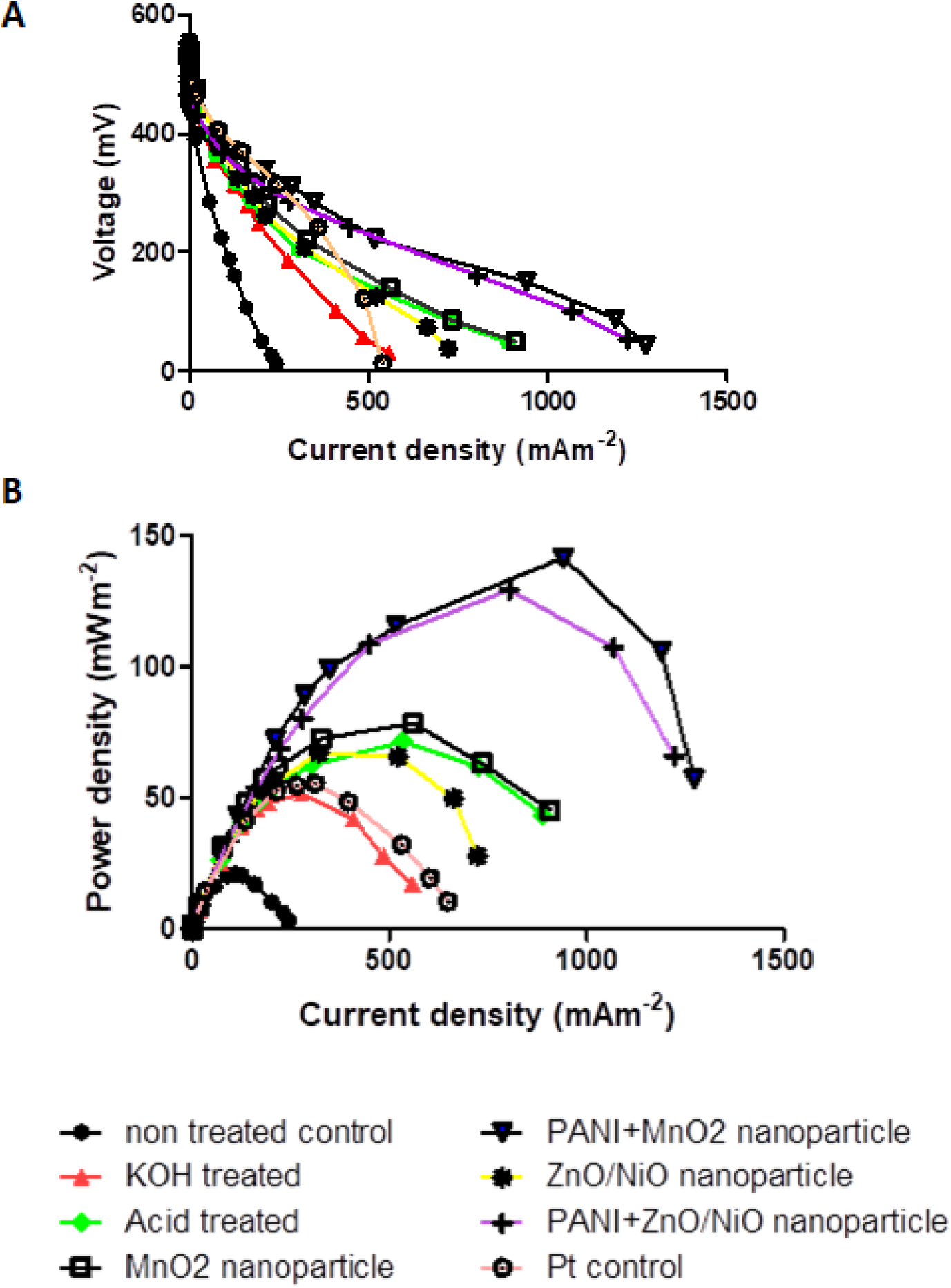
(A) Polarization plots of MFCs with different treated Carbon microfiber material electrodes as cathode and (B) Power-Current plots of MFCs with different, treated Carbon microfiber material electrodes as cathode.

**Table 1:** Maximum current density (J_max_), maximum power density (P_max_), and open circuit voltages recorded in MFCs with each treated carbon microfiber material electrode used as the cathode.

| Treatment | Maximum current density (mA/m<sup>2</sup>) | Maximum Power density (mW/m<sup>2</sup>) | Maximum open circuit voltage (mV) |
| --- | --- | --- | --- |
| Non treated Cathode | 244.30 ± 72.78 | 20.96 ± 6.54 | 509.3 ± 90.07 |
| KOH treated Cathode | 557.58 ± 106.58 | 51.7 ± 8.34 | 513.4 ± 37.28 |
| Acid treated Cathode | 889.7 ± 158.5 | 71.4 ± 6.4 | 503.1 ± 21.5 |
| Zinc treated Cathode | 724.1 ± 45.47 | 67.6 ± 1.50 | 573.8 ± 24.72 |
| Zinc oxide+ Nickel oxide + Polyaniline treated Cathode | 1222.7 ± 196.56 | 129.4 ± 7.34 | 502.3 ± 12.57 |
| Manganese treated Cathode | 908.6 ± 172.16 | 78.5 ± 5.08 | 538.7 ± 27.78 |
| Manganese oxide+ Polyaniline treated Cathode | 1272.7 ± 156.93 | 141.6 ± 22.06 | 456.9 ± 80.54 |
| Pt loaded control | 54.3 ± 12.2) | 642.1 ± 39.3 | 497.2 ± 44 |

The polarization curves and power-current plots of the MFC experiments clearly demonstrate that the nanoparticle loading on carbon electrodes makes a significan improvement on the electrochemical performance of all MFCs tested. In particular, PANI+MnO2 and PANI+ZnO/NiO nanoparticle loaded carbon microfiber electrodes demonstrated a significant enhancement of their electrochemical performances over the non-treated carbon control electrodes, Pt loaded control electrodes and KOH/acid treated electrode containing MFC systems (Fig 3A and 3B).

A direct comparison of the power and current densities of the control (untreated) electrode materials suggests that treatment of the electrode materials aids in increasing ORR catalysis. All other treatments tested showed significant increases in both power and current densities when compared to the control (Figure 4). As a result, MFC with MnO_2_/PANI treated electrode achieved the highest power density of 141.6 ± 22.06 mW/m^2^, which is 7-fold higher than the control (20.96 ± 6.54 mW/m^2^) with no treatment. PANI+MnO2 and PANI+ZnO/NiO nanoparticle incorporated electrodes also demonstrated more than 2-fold increases in power and current densities when compared to Pt loaded control MFC experiments.

**Figure 4:**
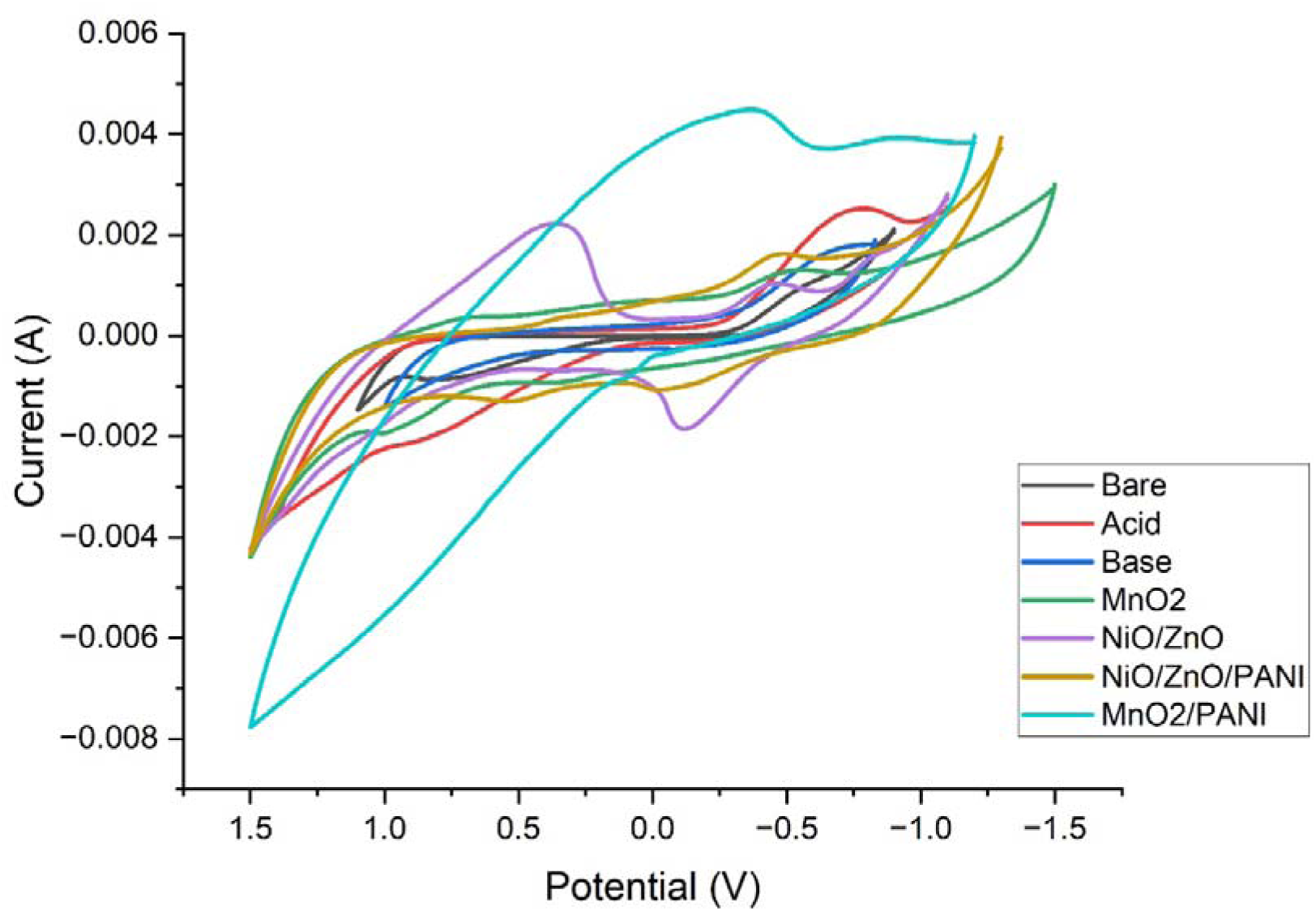
Cyclic voltammograms acquired at a scan rate of 50 mV/s on bare CMM and all other chemically functionalized and nanoparticles deposited CMM electrodes.

The findings of this study demonstrated that notably high-power densities can be obtained using metal oxides/polyaniline nanoparticles incorporated woven carbon microfiber cathodes devoid of noble metal catalysts such as platinum powder.

### 3.2 Cyclic voltammetry analysis of the performance of electrodes

Cyclic voltammetry is an electrochemical technique that has been widely used to describe the electron transfer interactions in the interface electrode-biomass/biofilm-liquor in the anode chamber or in the cathode/catholyte interface of MFCs.

CV involves scanning the voltage of a working electrode forward and backward at a predetermined pace (often 5 to 100 mV s^-1^). The reversible nature of the forward and reverse sweep plots of cyclic voltammograms represent a reversible oxidation and reduction reaction within the electrochemical system tested and therefore, indicative of the presence of reversible electrochemical reactions taking place within it. The presence of any mediators is detected by the production of a current during the mediator’s cyclic oxidation and reduction if the mediator can take (accept) reducing equivalents from the working electrode. The current-voltage map may then be used to calculate the mediator’s midpoint potential (Cheng et al., 2009).

A comparison of cyclic voltammograms acquired at a scan rate of 50 mV/s on bare CMM and all other treated electrodes used in this study is indicative of the presence of large oxidation and reduction CV peaks in the nanoparticle embedded MFC cathodes (Fig.4).

Generally, in cyclic voltammetry (CV), higher current values on the Y-axis during both forward and reverse sweeps indicate that oxidation/reduction reactions occur more readily for the tested material. Based on the obtained CV plots, it was observed that MnO_2_/PANI (manganese dioxide/polyaniline) exhibits more readily reversible electron transfer for oxygen reduction reaction (ORR), making it the most effective catalyst material among the modifications tested in the MFC system. The CV plots of other treatments also showed reversible oxidation/reduction peaks, indicating that none of the materials were irreversibly oxidized.

### 3.3 Scanning electron microscopy (SEM) analysis of functionalized microfiber electrode surfaces

Substrate selection is critical in the growth of nanomaterials such as stainless steel, nickel foam, and carbon cloth, which are commonly used as substrates for electrode material deposition in the case of nanomaterial electrode treatments. We chose woven carbon microfiber material (CMM) as the substrate in this experiment to increase the efficiency of the electron transfer due to high surface area, cost effectiveness. It was deemed a good substrate for nanomaterial deposition because of its large surface area and meshwork structure. This may allow the deposited nanomaterials to participate in electrochemical reactions and catalysis.

A morphological investigation is one of the available means to understand the prepared electrode samples. Supplementary fig. S5 shows SEM images of each treatment before and after they have been introduced to the MFC at the magnification of 1000x. It is evident from these images that the directly synthesize nanoparticle layer in each case of the electrode modification is robust enough to withstand the shear forces of air sparging into the MFC catholyte. This is indicative of sustained maintainence of good ORR catalysis levels through the application of the nanoparticle direct synthesis onto the CMM materials in MFC cathodes.

From the SEM images it is evident that the incorporated nanoparticles have remained on the novel microfiber electrodes and have been essential in catalyzing the ORR in the cathode compartments of the tested MFC systems.

Figure 5 provides a surface morphological comparison between the nanoparticles deposited CMM electrodes versus the bare CMM electrodes (fig 5 a, b and c compared to d, e and f). It indicates that almost the entire available surface area of the CMM fibrils are fully occupied by nanomaterial types directly synthesized onto the CMM surface. This suggests a good surface area occupied by the directly synthesized nanomaterials to conduct the ORR more efficiently. This could account for the superior electrochemical performance these directly synthesized nanoparticles produce compared to the traditional Pt powder catalyst loading onto CMM electrode surfaces using a conductive binder solution such as Nafion resin, where the distribution and the attachment of the catalyst material on the CMM surfaces is expected to be sparse.

**Figure 5:**
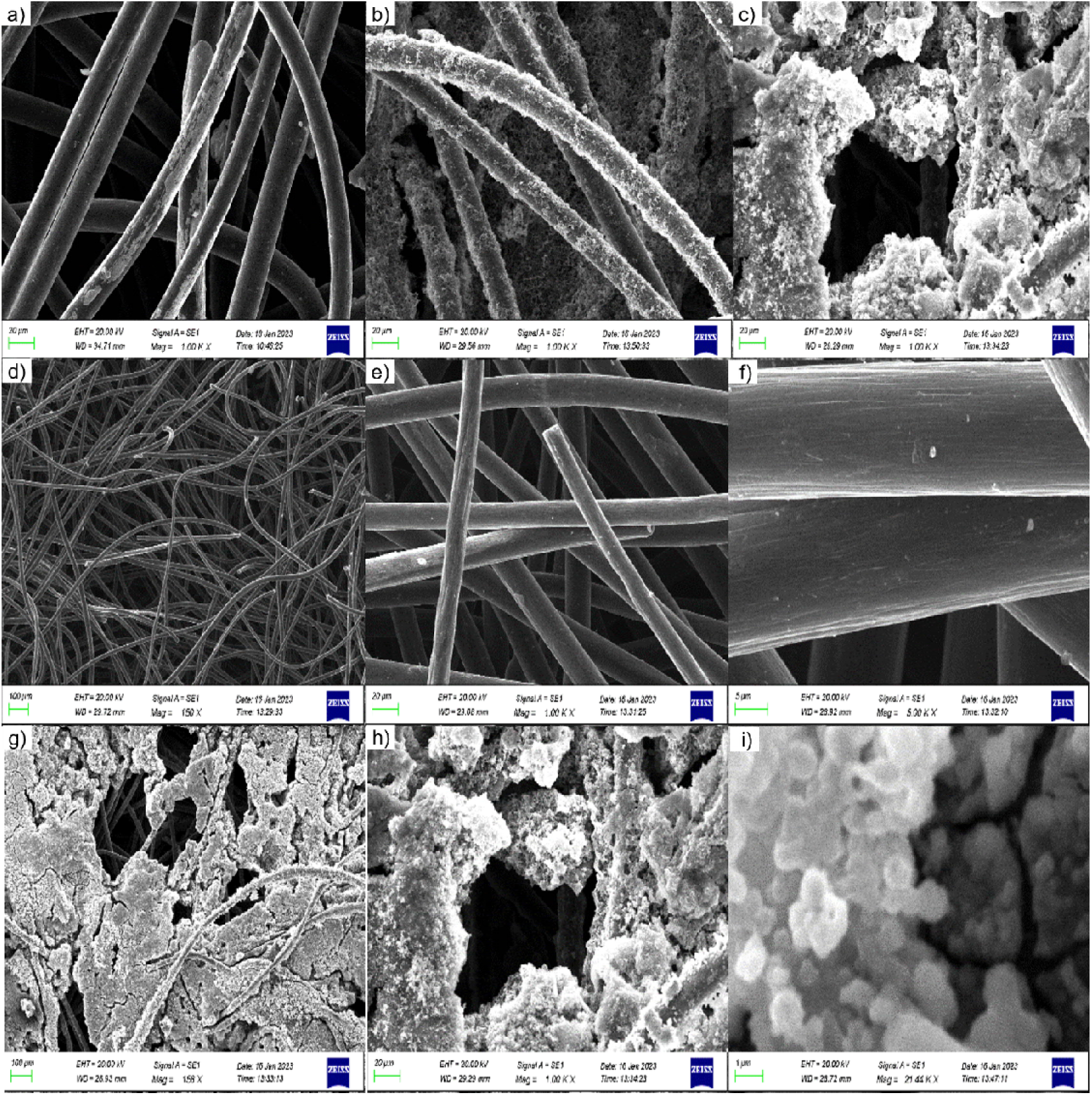
(a) MnO2 is directly deposited on the electrode material. (b) PANI synthesized by the electrodeposition method on the electrode (c) MnO_2_/PANI nanomaterials are deposited on the electrode material. (d) Bare Carbon microfiber material under 150x magnification (e) 1000x magnification (f) 5000x magnification. (g) MnO_2_ and PANI treated Carbon microfiber material under 150x magnification (h) 1000x magnification (i) 22.44k x magnification.

The CMM as the substrate has provided a well-connected porous platform for uniform and wide distribution of PANI on it. This design aided in the reduction of fabrication time for electrodes. The PANI on the carbon electrode fiber network seemed to be well distributed and adhered to the CMM that could exhibit good conductivity and stability. Additionally, the electrodes can be designed binder-free as PANI is directly deposited on the CMM. The neatly deposited PANI with CMM can be an efficient electrode for MFC applications.

The elemental analysis and mappings of the electrodes were obtained by Energy Dispersive X-Ray Analysis (EDAX) spectra. Fig. 6 indicates EDAX spectra of Non-treated CMM, MnO_2_/CMM, and MnO_2_/PANI/CMM electrodes before and after introducing them to MFCs under 150x magnification (a and b). Figure 6 (c) shows the EDAX spectrum of MnO_2_/CMM before introducing it to the cell, where the respective peaks corresponding to the elements carbon, nitrogen, Oxygen, and Manganese are detected in the treated electrode surface with a 150X magnification. Figure 6 (d) shows the MnO_2_/CMM electrode after being introduced to the cell, where the respective peaks corresponding to the elements carbon, nitrogen, Oxygen, Phosphorus, and Potassium and a small amount of Manganese are detected in the treated electrode surface with a 150X magnification. Phosphorus and Potassium peaks, appearing in the EDAX spectrum of the MnO_2_/CMM surface after being introduced to the cell, are due to the Phosphate buffer used as the media of the cathode chamber.

**Figure 6:**
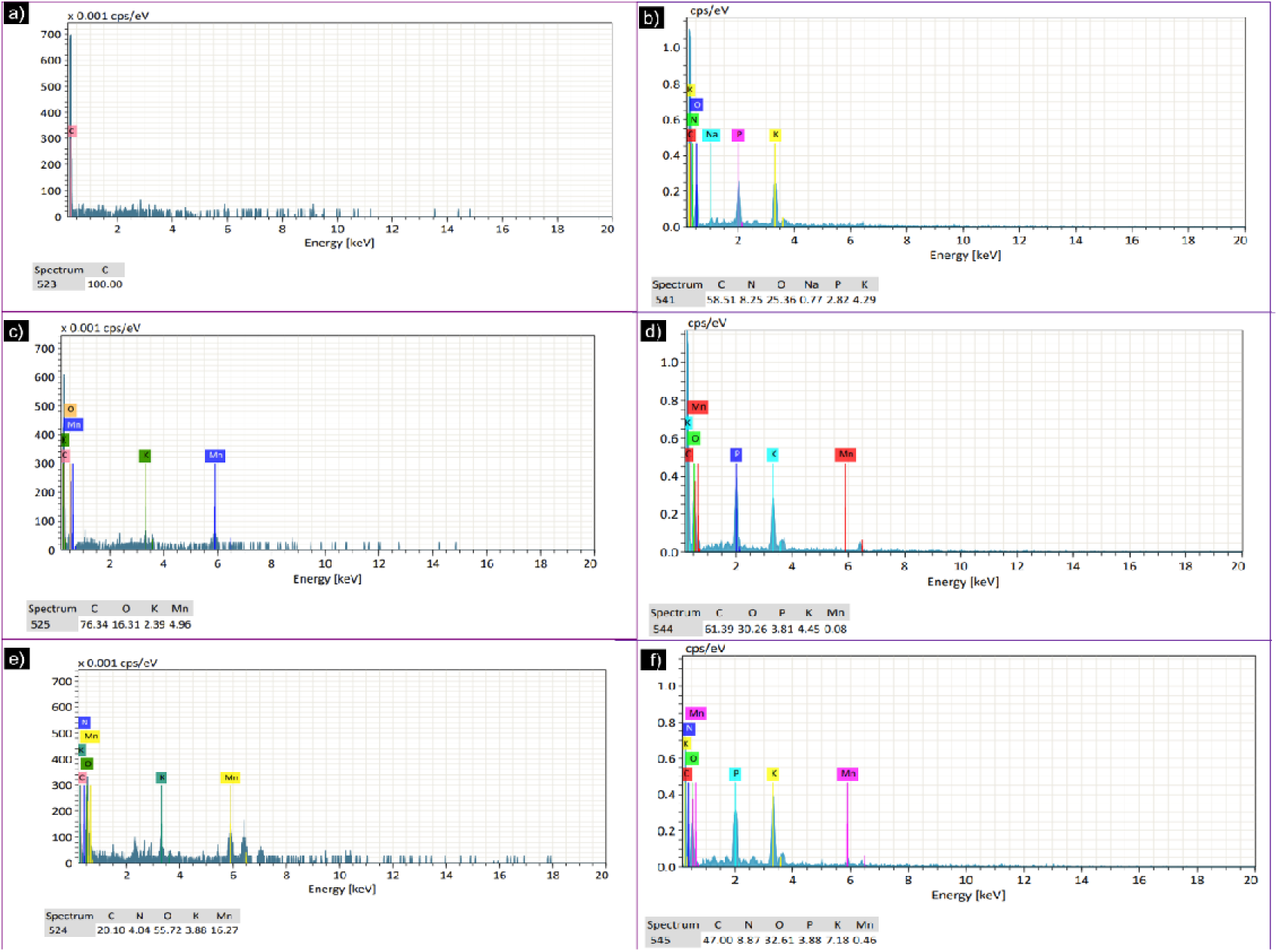
Elemental analysis and mapping of bare CMM electrodes and nanoparticle modified CMM electrodes obtained using Energy Dispersive X-Ray analysis under magnification of 150x.

The elemental mapping images, shown in Figures 6 (e) and (f), reveal the uniform spatial distribution of the elements present in the MnO_2_/PANI/CMM before introducing the electrode to the cell and after introducing the electrode to the microbial fuel cell.

### 3.4 Fourier Transform Infrared Spectroscopy (FTIR) analysis of functionalized cathodes

FTIR can provide information about the presence of functional groups on the surface of nanomaterials. By analyzing the infrared absorption spectra, it is possible to identify the types of chemical bonds and functional groups present in the nanomaterials, which can help in understanding the surface chemistry and composition of the modified electrodes that may ultimately improve the ORR performance of the novel electrodes. FTIR can be used to study the changes in the surface chemistry of the electrodes after nanomaterial modification. By comparing the FTIR spectra of the unmodified and modified electrodes, it is possible to determine the specific chemical changes that occur on the electrode surface as a result of nanomaterial deposition or surface functionalization.

It was confirmed by FTIR analysis that the woven CMM electrodes were modified with nanomaterials following treatment with metals and metal oxides.

Figure 10 compares the FTIR spectra of bare and treated electrodes.

The nanomaterial-treated CMM electrodes indicate the emergence of many additional major peaks compared to the Non treated electrodes.

The FTIR spectra (Supplementary fig. S1) of treated electrodes indicate key spectral differences between the nanoparticle treated electrodes and non-treated control electrodes in MFC cathodes

The obtained FTIR spectra of MnO_2_ and MnO_2_/PANI show the broad wave number bands at 3347.7cm^-1^ correspond to N-H stretching. The wave number band at 1739.38 cm^-1^ shows a C=O which may represent the carbon and oxygen bond system of the carbon microfiber electrodes. Among the various peaks assigned to PANI, the characteristic peaks around 1654.12 cm^-1^ correspond to benzenoid (-(C_6_H_4_)-) ring, C-H bond. Another main band at 1394.75 cm^-1^ can be assigned to the stretching of C-N in -NH-(C_6_H_4_)-NH-. The bands appeared at 876.87 cm^-1^ which corresponds to the stretching of C-H bonds.

## 4.0 Conclusion

The findings of this study demonstrated that using nanomaterials-modified carbon microfiber electrode material can significantly improve the electrochemical performance of lake sediment inoculated microbial fuel cells. This can be used as an alternative to expensive cathode catalysts like platinum in MFCs to catalyze ORR. The electrochemical performances obtained in this study by using these nanomaterial-modified electrodes in two-chamber MFCs were comparable to or better than the P_max_ and J_max_ values obtained in previous studies by operating similar two-chamber MFC systems employing noble metals as the ORR catalysts. In particular, the MnO_2_ + PANI incorporated cathodes and ZnO/NiO + PANI nanoparticle incorporated cathodes demonstrated approximately 6 fold increase in P_max_ and J_max_ values over non-treated cathode controls.

## Supporting information

Supplementary data - main

## Acknowledgments

Technical staff who assisted in carrying out the experimental work in numerous ways at all collaborating institutions and laboratories are thankfully acknowledged. Authors wish to acknowledge the Rajarata University of Sri Lanka (RUSL) for providing funding, materials and instrumentation support under the microbiology special degree program for conducting and completing this research. They also wish to thank the Faculty of Technology, RUSL for providing access to SEM and EDX analysis of specimens.

