## Supplementary data - main for "Use of nanomaterials-modified carbon microfibre electrode material for superior electrochemical performance in lake sediment inoculated microbial fuel cells"

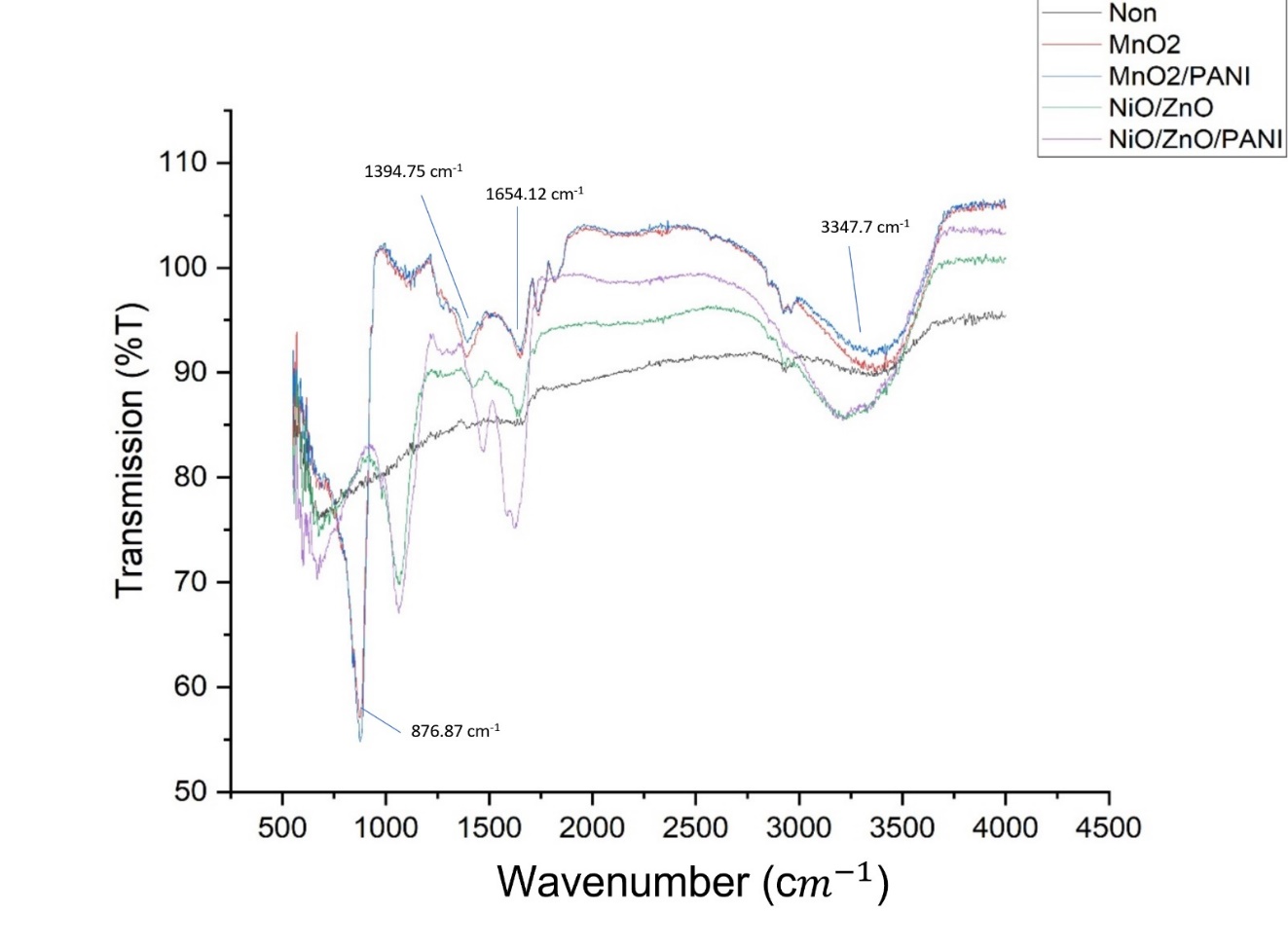

**Supplementary Figure S1:** The FTIR spectra of treated electrodes indicate key spectral differences between the nanoparticle treated electrodes and non-treated control electrodes in MFC cathodes

### **Isolation of anodic microorganisms**

This medium is used to isolate microorganisms from the anode. After two to three days of microorganism growth in the media inside an anaerobic jar (Supplementary Figure 2) the media had turned colorless, and the suspended black powder was not visible (Supplementary Figure 3).

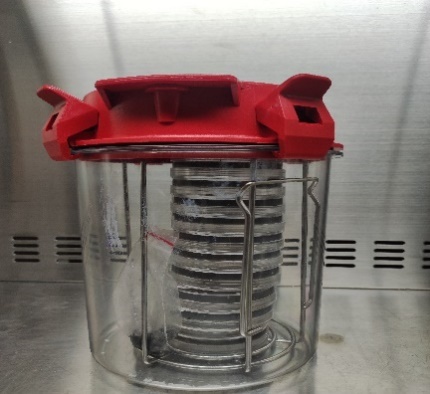

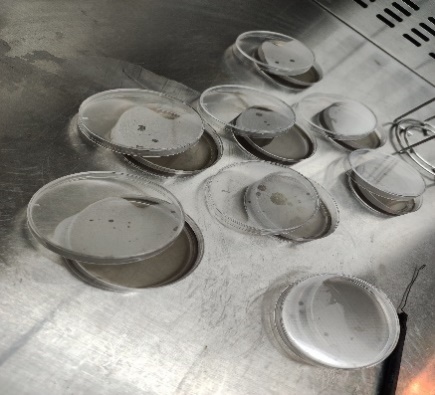

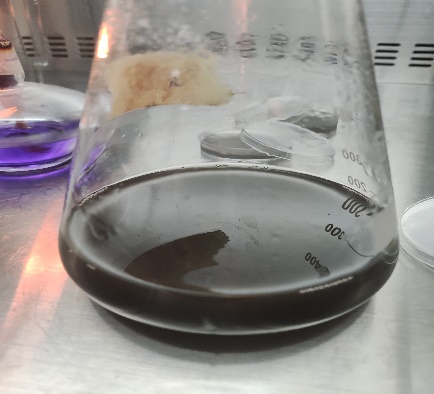

(b)

(c)

(a)

**Supplementary Figure S2**: Isolation of anodic micro-organisms (a) novel chromogenic media with MnO_2_ particles (b) Petri dishes containing chromogenic media (c) Growth condition for anaerobic microorganisms inside an anaerobic jar

| 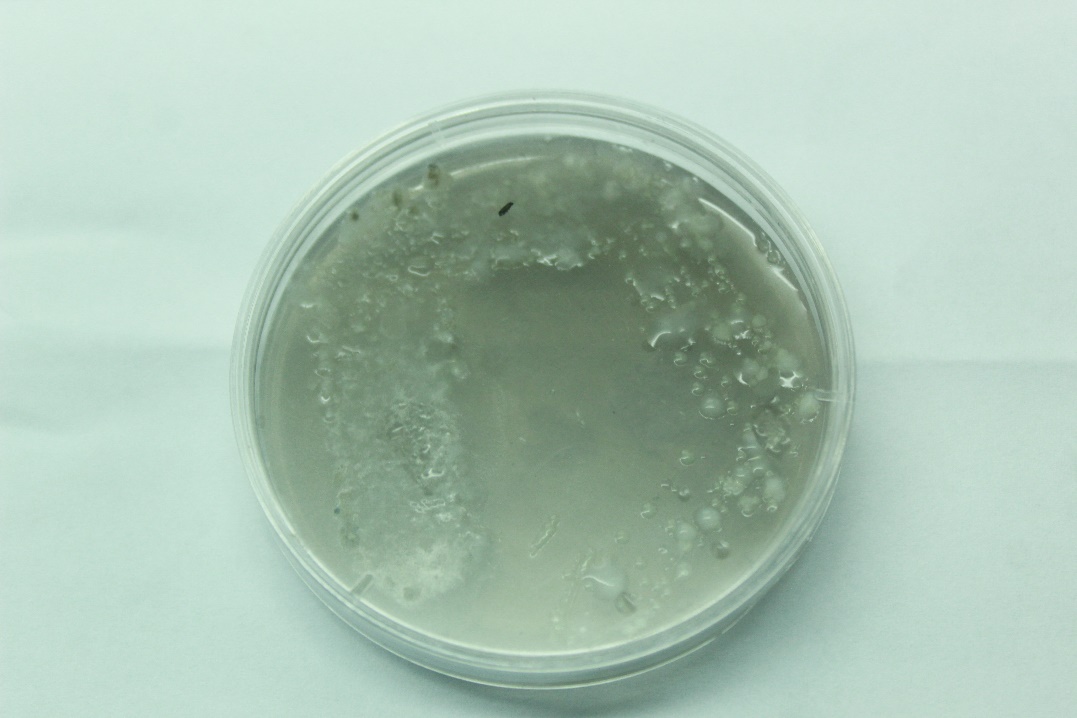 | 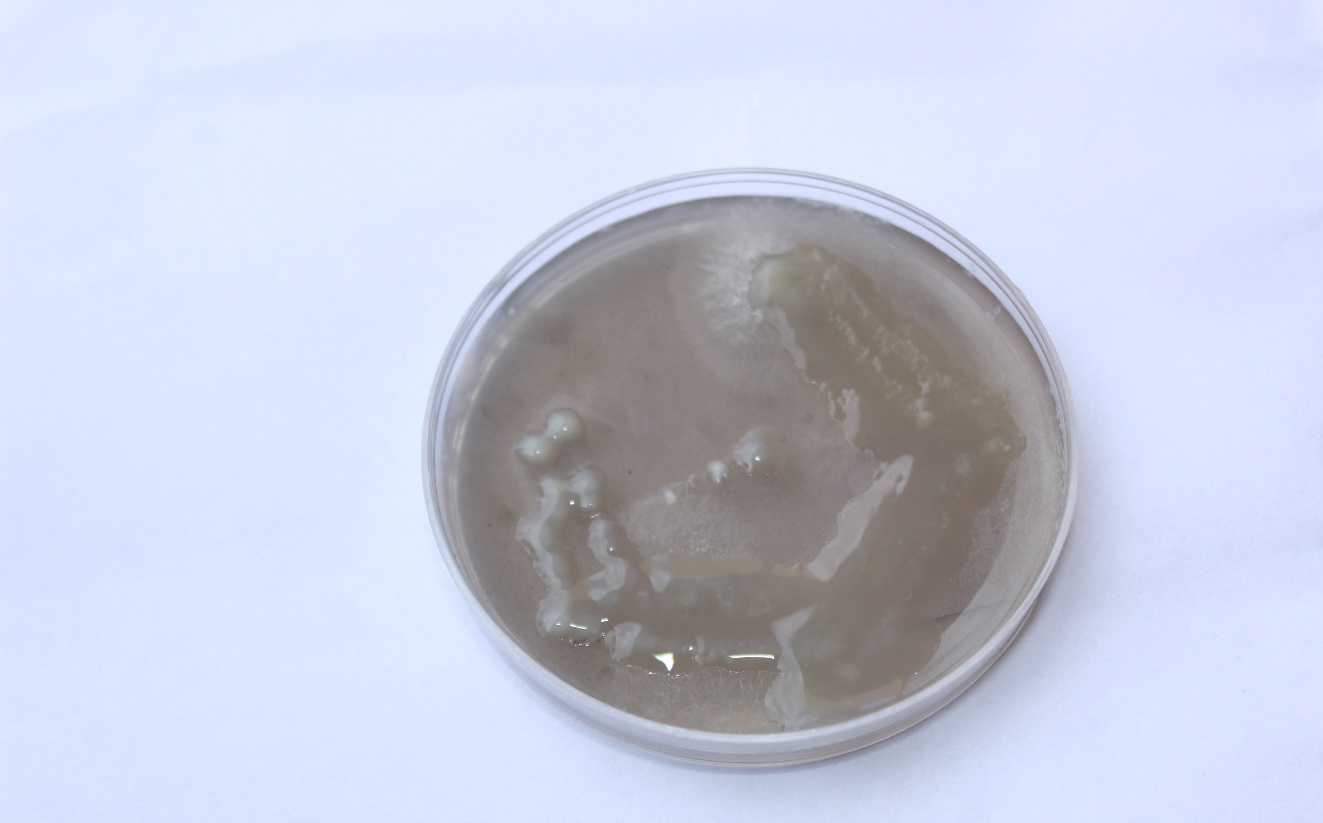 |
| --- | --- |

**Supplementary Figure S3**: Exoelectrogenic microbes isolated from the anode compartment.

### **Molecular identification of the anodic microorganisms**

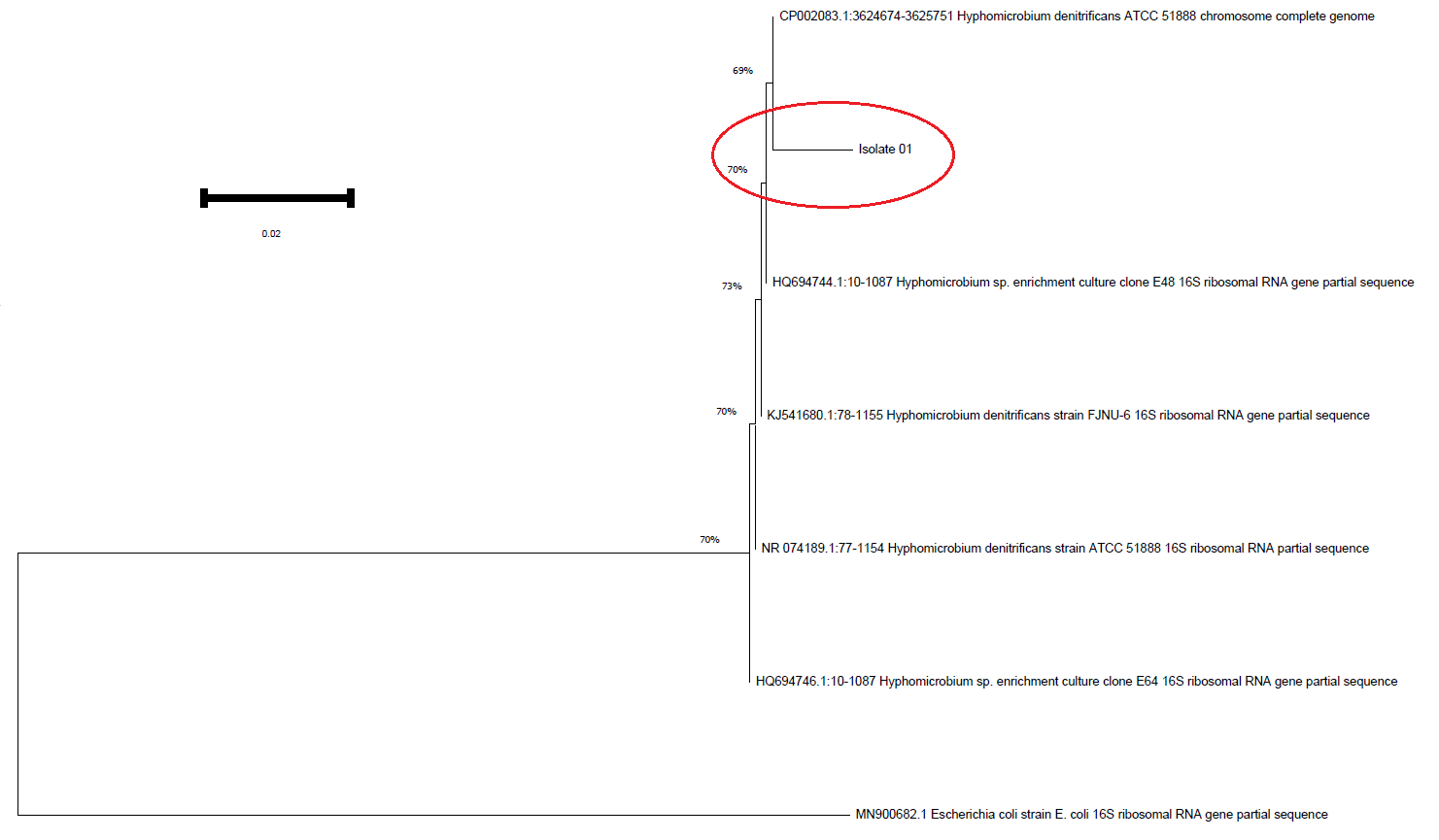

**Supplementary Figure S4:** The evolutionary history was inferred using the Neighbor-Joining method. The optimal tree is shown. The evolutionary distances were computed using the Maximum Composite Likelihood method and are in the units of the number of base substitutions per site. The proportion of sites where at least 1 unambiguous base is present in at least 1 sequence for each descendent clade is shown next to each internal node in the tree. This analysis involved 7 nucleotide sequences. All ambiguous positions were removed for each sequence pair (pairwise deletion option). There were a total of 1545 positions in the final dataset. Evolutionary analyses were conducted in MEGA11.

According to the Neighbor-Joining tree obtained the Isolate 01 shows 69% similarity to *Hyphomicrobium denitrificans.*

*Hyphomicrobium denitrificans* is a bacterial species in the Hyphomicrobiaceae family. It is a Gram-negative, rod-shaped bacterium found in a variety of aquatic environments, including freshwater and marine environments. This bacterium is mostly heterotrophic, which means it gets its energy and carbon from organic compounds. It can grow on a variety of organic substrates, including sugars, alcohols, organic acids, and amino acids.

*Hyphomicrobium denitrificans* can be found in a variety of aquatic environments, including rivers, lakes, sediments, and wastewater treatment systems. It is also possible to isolate it from soil and biofilms.

**Isolation of anodic microorganisms**

Chromogenic and media are microbiological growth media containing enzyme substrates linked to a chromogen (color reaction), a fluorogenic (light reaction), or both. MnO2, an insoluble black crystalline powder suspended in the growth media used here, gives the media an overall black color (Supplementary figure 2).

Manganese (Mn) is in its +4-oxidation state in MnO2, resulting in a black, insoluble powder. This finding shows that these anodic microorganisms can convert MnO2 into more soluble and colorless Mn2+ ions. This demonstrates the ability of anodic bacteria to act as exoelectrogenic bacteria. Because MNO2 is insoluble, it cannot be taken into the cell and used by the cell; thus, the only way to reduce MnO2 particles is to transfer electrons from the cell to the MNO2 granules (Nazeer Z. & Fernando, 2022). As a result, the microbes isolated using this selective medium are exoelectrogens capable of using MNO2, an insoluble electron sink as a terminal electron acceptor in their metabolic activities. The identity of the isolated exoelectrogenic bacteria from the MFC was established using 16s rRNA amplicon sequencing. The identity of the isolated exoelectrogen was established as Hyphomicrobium denitrificans, when placed in a phylogenetic tree (Supplementary fig. S4)

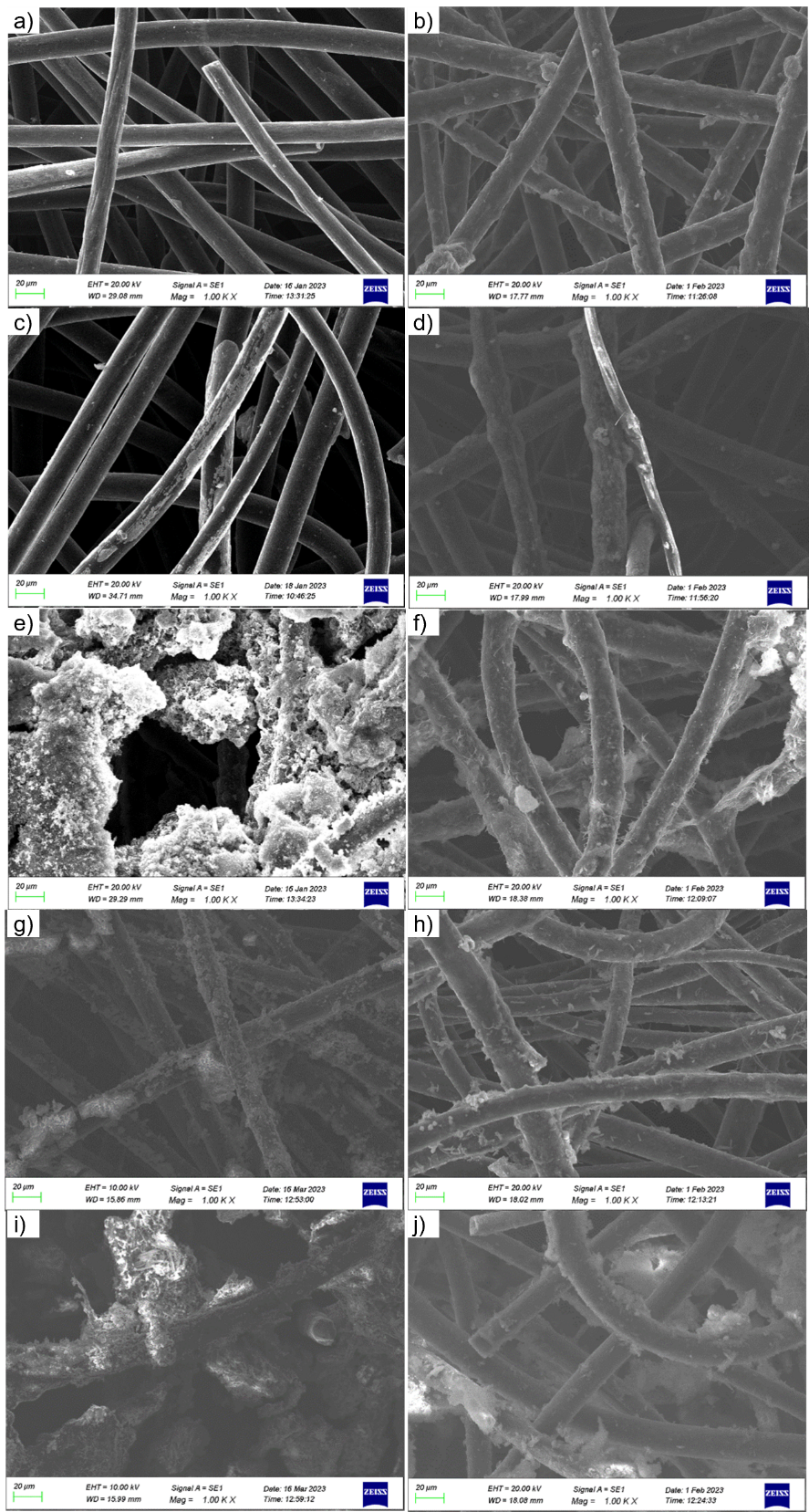

**Supplementary Figure S5:** SEM images of the nanoparticles incorporated electrodes before and after they have been introduced into the MFC cathode with 1000x magnification. a) Non-treated electrode before introducing to the MFC b) Non-treated electrode after introducing to the MFC c) MnO_2_/CMM electrode before introducing to the MFC d) MnO_2_/CMM electrode after introducing to the MFC e) MnO_2_/PANI/CMM electrode before introducing to the MFC f) MnO_2_/PANI/CMM electrode after introducing to the MFC g) ZnO/NiO/CMM electrode before introducing to the MFC h) ZnO/NiO/PANI/CMM electrode after introducing to the MFC

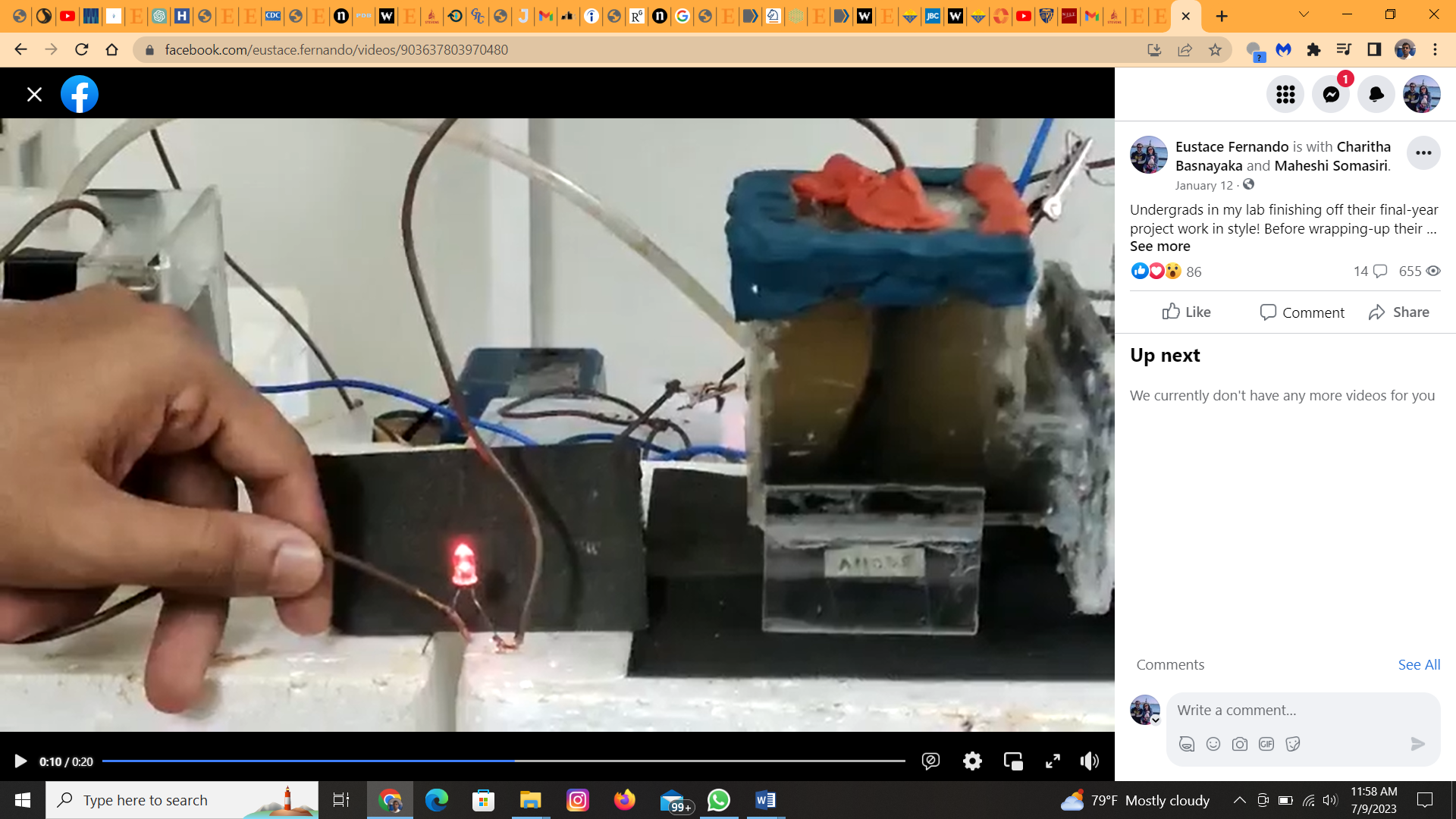

**Supplementary figure S6:** MnO_2_/PANI/CMM electrode incorporated MFCs being used to power small electronic devices such as light emitting diodes (LEDs) in this demonstration indicated that nanomaterials modified electrodes produce sufficient current and voltage to power small electronic devices.
